# Tcf21 as a Founder Transcription Factor in Specifying Foxd1 Cells to the Juxtaglomerular Cell Lineage

**DOI:** 10.1101/2024.03.25.586641

**Authors:** Hina Anjum, Jason P. Smith, Alexandre G Martini, George S. Yacu, Silvia Medrano, R. Ariel Gomez, Maria Luisa S. Sequeira-Lopez, Susan E. Quaggin, Gal Finer

## Abstract

Renin is crucial for blood pressure regulation and electrolyte balance, and its expressing cells arise from Foxd1+ stromal progenitors. However, factors guiding these progenitors toward renin-secreting cell fate remain unclear. Tcf21, a basic helix-loop-helix (bHLH) transcription factor, is essential in kidney development. Utilizing *Foxd1*^*Cre/+*^*;Tcf21*^*f/f*^ and *Ren1*^*dCre/+*^*;Tcf21*^*f/f*^ mouse models, we investigated the role of Tcf21 in the differentiation of Foxd1+ progenitor cells into juxtaglomerular (JG) cells. Immunostaining and in-situ hybridization demonstrated fewer renin-positive areas and altered renal arterial morphology, including the afferent arteriole, in *Foxd1*^*Cre/+*^*;Tcf21*^*f/f*^ kidneys compared to controls, indicating Tcf21’s critical role in the emergence of renin-expressing cells. However, Tcf21 inactivation in renin-expressing cells (*Ren1*^*dCre/+*^*;Tcf21*^*f/f*^) did not recapitulate this phenotype, suggesting Tcf21 is dispensable once renin cell identity is established.

Using an integrated analysis of single-cell RNA sequencing (scRNA-seq) and single-cell assay for transposase-accessible chromatin sequencing (scATAC-seq) on GFP+ cells (stromal lineage) from E12, E18, P5, and P30 *Foxd1*^*Cre/+*^*;Rosa26*^*mTmG*^ control kidneys, we analyzed the temporal dynamics of Tcf21 expression in cells comprising the JG lineage (*n*=2,054). A pseudotime trajectory analysis revealed that Tcf21 expression is highest in metanephric mesenchyme and stromal cells at early developmental stages (E12), with a decline in expression as cells mature into renin-expressing JG cells. Motif enrichment analyses supported Tcf21’s significant involvement in early kidney development. These findings underscore the critical role of Tcf21 in Foxd1+ cell differentiation into JG cells during early stages of kidney development, offering insights into the molecular mechanisms governing JG cell differentiation and highlight Tcf21’s pivotal role in kidney development.

**NEW & NOTEWORTHY:** This manuscript provides novel insights into the role of Tcf21 in the differentiation of Foxd1+ cells into JG cells. Utilizing integrated scRNA-seq and scATAC-seq, the study reveals that Tcf21 expression is crucial during early embryonic stages, with its peak at embryonic day 12. The findings demonstrate that inactivation of Tcf21 leads to fewer renin-positive areas and altered renal arterial morphology, underscoring the importance of Tcf21 in the specification of renin-expressing JG cells and kidney development.

## INTRODUCTION

Juxtaglomerular (JG) cells are specialized smooth muscle cells located in the walls of the kidney’s afferent arterioles, playing a crucial role in regulating blood pressure and fluid balance through the secretion of renin (1-3). The differentiation of these cells from their progenitors is a tightly regulated process, involving various transcription factors (4-6). Renin cells originate during kidney development from the stromal progenitors, the Forkhead box D1 positive (Foxd1+) cells, within the metanephric mesenchyme, around embryonic day 13.5 (E13.5) of the mouse (7). Tcf21, a member of the basic helix-loop-helix (bHLH) transcription factor family, has been implicated in the development of several mesenchymal cell lineages, including those in the kidney (8, 9). However, its specific role in the differentiation of Foxd1+ progenitor cells into JG cells remains unclear. Foxd1 is a transcription factor critical for the formation and differentiation of renal stromal cells, which give rise to several cell types, including JG cells (10-12). Previous studies have demonstrated that Foxd1+ progenitors are essential for kidney morphogenesis and vascular development (10). Given the known roles of Tcf21 and Foxd1 in kidney development, it is hypothesized that Tcf21 may play a pivotal role in the differentiation of Foxd1+ cells into JG cells. This study aims to elucidate the role of Tcf21 in the specification and differentiation of Foxd1+ progenitor cells into renin-expressing JG cells. Utilizing *Foxd1*^*Cre/+*^*;Tcf21*^*f/f*^ and *Ren1*^*dCre/+*^*;Tcf21*^*f/f*^ mouse models, histological analysis, and integrated scRNA-seq and scATAC-seq analysis, we investigated the temporal and spatial dynamics of Tcf21 expression in these cells across various developmental stages. Our results indicate that Tcf21 is crucial for the early specification of Foxd1+ cells into the JG lineage, with its expression peaking during the early embryonic stage, and decline as kidney development progresses. This study provides new insights into the molecular mechanisms governing JG cell differentiation and highlights the importance of Tcf21 in kidney development.

## MATERIALS AND METHODS

### Mice and genotyping

All mouse experiments were approved by the Animal Care Committee at the Center for Comparative Medicine of Northwestern University (Evanston, IL) and were performed in accordance with institutional guidelines and the NIH Guide for the Care and Use of Laboratory Animals. The *Tcf21*^*flox/flox*^ and *Ren1*^*dCre/+*^ mice were created as previously described (13, 14). The *Foxd1-eGFPCre* mice were kind gifts from Dr. Andrew McMahon (University of Southern California) and are described elsewhere (15). Genotyping was performed with Quick-Load Taq 2X Master Mix (M0271L, BioLabs), on a 2% agarose gel for the PCR bands. Primer information is in **Suppl. Table 1**.

### Histology, immunohistochemistry and immunofluorescence

Mouse embryo dissections were performed at the indicated time-points. Samples were fixed with 10% Neutral buffered formalin, 4% paraformaldehyde, or Bouin’s solution overnight, embedded in paraffin, and sectioned into 4 µm by the Mouse Histology and Phenotyping Laboratory (MHPL) of Northwestern University. Sections were deparaffinized in xylene and ethanol and boiled in citrate buffer (10 mM sodium citrate and 0.5% Tween-20, pH = 6.0) for antigen retrieval. Sections were then washed with 1% BSA in PBS, permeabilized with 0.3% Triton-X-100, and blocked with 5% donkey serum and 0.3% Triton-X-100 in PBS before incubating with primary antibodies at 4°C overnight. Sections were then washed in PBST and incubated with Alexa Fluor fluorescence-conjugated secondary antibodies for 90 minutes at room temperature. Antibody information: CD31 (R&D, AF3628, 1:100), CD146 (abcam, ab75769, 1:200), Krt8 (TROMA-I-s, DSHB. 1:20), Renin (Alpha Diagnostic, Gomez project 1:500). All secondary antibodies were used at a 1:200 dilution.

### In-situ hybridization (RNAScope)

In-situ hybridization was performed on paraffin section using RNAscope (16) and following ACD^®^ protocol https://acdbio.com/rnascope-25-hd-duplex-assay. Probes: Tcf21 (ACD #508661), Ren1 (ACD #433461).

### Image quantification

MCAM+ Cells (E16.5 and E18.5 Kidneys): Confocal fluorescent images of interlobular arteries and surrounding glomeruli were acquired with four z-stacks at 0.75-µm intervals using a 40X objective. Images were pre-processed with Gaussian filtering (0.25-µm radius) to reduce noise. MCAM+ areas were segmented using the Li thresholding algorithm in FIJI (ImageJ, v2.14.0). Interlobular arteries were identified by MCAM+ staining on the inner layer and CD31+ staining on the outer layer. The MCAM+ area for each artery was measured across z-slices, and the average area was used for quantification.

Vessel Density, Branching Index, and Vessel Length (E14.5 and E18.5 Kidneys): Confocal fluorescent images were acquired of whole kidneys at E14.5 or kidney sixths at E18.5, with each sixth analyzed individually. Non-kidney portions of the images were removed in Adobe Photoshop and CD31+ levels were normalized across all images. CD31+ signals were then analyzed using AngioTool software, with parameters set for vessel diameter (10 µm), thresholding (15-255), and exclusion of small particles below 400 µm^2^.

Renin+ Regions in Glomeruli (E18.5 Kidneys): Confocal fluorescent images of all Renin+ regions within the kidney were acquired using a 40X objective. Images were pre-processed with Gaussian filtering (0.25-µm radius) and segmented using the Max Entropy thresholding algorithm in FIJI. Each Renin+ region was classified by a trained investigator into two categories: glomerular or arterial based on their localization with the juxtaglomerular apparatus or renal arterial wall. The number of glomeruli with Renin+ regions was counted and normalized to the total number of glomeruli for each kidney. Additionally, the area of Renin+ regions in glomeruli was measured and normalized to the glomerular area. To measure the glomerular area, glomeruli were manually identified by CD31 signal with a 25X objective. Ellipses of best fit were drawn around each glomerulus and the areas were summed for each kidney.

### Isolation of kidney single cells: E12

Pregnant mice were anesthetized with tribromoethanol. Small fetuses were removed, and GFP pups were identified using the EvosFLC cell imaging system (LifeTechnologies, California, USA). The metanephros area was dissected, minced, and incubated in TryPLE Express (Gibco, New York, USA) at 37°C for 5 minutes. After adding DMEM + 5% FBS, the mixture was homogenized and filtered through a 40 µm nylon cell strainer. The filtrate was centrifuged, and the pellet resuspended in resuspension buffer, preparing cells for single-cell capture. n=15 (E12, both sexes represented based on Mendelian ratios).

### Isolation of Kidney Single Cells: E18, P5, and P30

Animals were anesthetized with tribromoethanol (300 mg/kg). Kidneys from P5 and P30 mice were excised, decapsulated, and dissected. The kidney cortices were minced and digested in an enzymatic solution (0.3% collagenase A, 0.25% trypsin, 0.0021% DNase I) in a shaking incubator (80 RPM) at 37°C for 15 minutes. The supernatant was collected, and the digestion was repeated three times. The pooled supernatants were centrifuged, and the pellet resuspended in buffer 1 (130 mM NaCl, 5 mM KCl, 2 mM CaCl2, 10 mM glucose, 20 mM sucrose, 10 mM HEPES, pH 7.4), filtered through 100 µm and 40 µm nylon cell strainers, and centrifuged again. The final pellet was resuspended in resuspension buffer, DAPI added, and GFP positive cells were sorted by FACS (Influx Cell Sorter or FACS Aria Fusion Cell Sorter, Becton Dickinson) and resuspended in DMEM with 10% FBS for immediate use. n=18 (E18, both sexes represented based on Mendelian ratios), n=12 (P5, 6 females and 6 males), and n=10 (P30, 4 females and 6 males).

### scATAC-seq Library Preparation

GFP-positive cells were washed in dPBS with 0.04% BSA and counted with a CellometerMini (Nexcelom Bioscience, Massachusetts, USA). Cells with over 80% viability were processed for nuclei isolation using Low Cell Input Nuclei Isolation protocol. Cells were lysed, washed, and nuclei counted with a Neubauer Chamber (Spencer, Buffalo, USA). Nuclei were captured using the Chromium System (10X Genomics) and processed for sequencing on an Illumina HiSeq platform.

### scRNA-seq Library Preparation

GFP-positive cells were spun down and resuspended in dPBS with 0.04% BSA, counted, and processed with the Chromium System (10X Genomics) following the manufacturer’s recommendations. Barcoded cDNA was amplified and sequenced on an Illumina HiSeq platform.

### Read Preprocessing

Genome and transcriptome annotations were performed using the mm10 genome. Cell Ranger pipeline (refdata-cellranger-arc-mm10-2020-A-2.0.0) and BSgenome package (BSgenome.Mmusculus.UCSC.mm10) were used.

### scATAC-seq Analysis

FASTQ files were processed with Cell Ranger ATAC (cellranger-atac-2.0.0), and duplicate reads were removed. ArchR v1.0.1 was used for quality control, dimensionality reduction, batch correction (Harmony algorithm), and cell clustering. Pseudo-bulk replicates were generated for peak calling using MACS2.

### scRNA-seq Analysis

FASTQ files were processed with Cell Ranger (cellranger-6.0.1). Feature-barcode matrices were analyzed with Seurat for normalization, PCA, clustering, and cell identification using marker genes. Integration of scRNA-seq and scATAC-seq data was performed using ArchR’s addGeneIntegrationMatrix function.

### JG-Cell Differentiation Trajectory

To construct a pseudotime trajectory of the differentiation of Foxd1+ stromal progenitors to mature renin-expressing cells, we first identified cell populations that would form the backbone of this trajectory using established markers of the beginning and end of this process. First, we identified all cells expressing *Foxd1* at or above the third quartile and which also displayed a predicted gene score based on chromatin accessibility in the upper quartile. Next, we identified cells with combined expression and chromatin accessibility based on gene score values in the upper quartile for both *Ren1* and *Akr1b7*. We established populations from early developmental timepoints and high *Foxd1* values as the start of the trajectory backbone, and cells during and late in development with high *Ren1/Akr1b7* values as the endpoints. Finally, we used the addTrajectory() and plotTrajectory() functions from ArchR to map the dynamic chromatin and gene expression changes.

## RESULTS

### Inactivation of *Tcf21* in Foxd1+ cells leads to fewer renin-positive areas in the developing kidney

Utilizing the *Foxd1*^*Cre/+*^*;Tcf21*^*f/f*^ mouse in which *Tcf21* is excised in Foxd1+ cells (17), we initially examined renin protein expression by immunostaining. At mid-and late-stages of kidney development (i.e., E16.5 and E18.5), *Tcf21* stromal conditional knockout (*Tcf21* stromal cKO) demonstrated globally fewer renin-positive areas at the juxtaglomerular apparatus and arteriolar walls compared with controls (**Fig.1 IA-C**). Morphologic examination of the developing afferent arterioles, where renin cells normally reside, revealed thinner and shorter walls in the E16.5 *Tcf21* stromal cKO compared with controls (**Fig.1 II**). To further corroborate this finding, we performed immunostaining for renin and the melanoma cell adhesion molecule (MCAM, CD146), a marker of perivascular cells (e.g. pericytes, VSMC) (18-20). This staining revealed reduced renin and MCAM expression in the walls of the afferent arterioles and radial arteries in *Tcf21* stromal cKO compared with control (**Fig.1 IIIA-C**). This renin+MCAM+ cell population has been previously characterized as renin-positive pericytes that are believed to represent a specialized subpopulation known for their pericyte role in supporting vessel formation and for their distinct ability to synthesize renin (2, 21).

**Figure 1:**
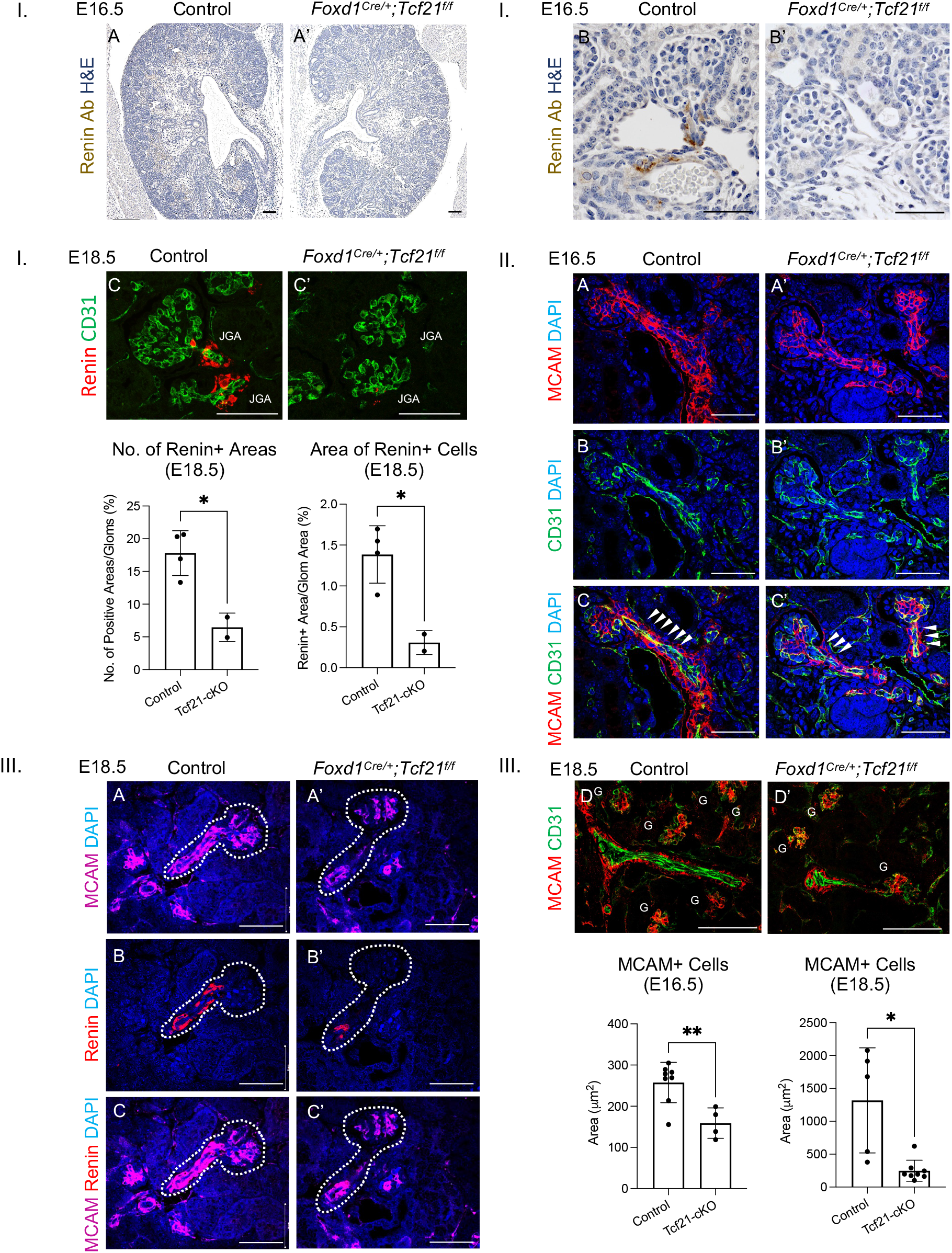
Inactivation of Tcf21 in Foxd1+ cells reduces renin expression at E16.5 and E18.5. **I**.A. Representative image of an E16.5 kidney from *Foxd1*^*Cre/+*^*;Tcf21*^*f/f*^ mouse showing decreased renin expression (brown staining) compared to control. *n*=4, sex not determined. B. Juxtaglomerular (JG) area of a glomerulus from a *Tcf21* stromal cKO demonstrates reduced renin expression compared to control (IHC on H&E staining). *n*=6. C. Top: Immunostaining with anti-renin antibody similarly shows reduced renin expression in the JG apparatus of *Tcf21* stromal cKO compared with control. Bottom: Quantifications of the number of renin-expressing cell aggregates (renin-positive areas) (left), and the area of renin-positive aggregates in controls and *Tcf21* stromal cKO kidneys at E18.5 (*p*=0.014 and *p*=0.016, respectively). *n=*6. **II**. A-C. Afferent arterioles stained for CD31+ (PECAM) endothelial cells and CD146+ (MCAM) pericytes are thinner and shorter in *Tcf21* stromal cKO compared to controls. *n*=12. **III**. A. reduced CD146+Renin+ staining along the afferent arteriole in *Tcf21* stromal cKO compared to control. *n*=4. B. Representative image of an interlobular artery from an E18.5 kidney in a *Foxd1*^*Cre/+*^*;Tcf21*^*f/f*^ mouse, showing a thinner and shorter compared to control (*G*=glomeruli). C. Quantifications of MCAM+ area (renin+ pericytes) at E16.5 (*p=*0.004) and E18.5 (*p*=0.039). *n*=4 (E16.5) and *n*=6 (E18.5). Sex not determined. Scale bar 50μm.

Complementarily, we evaluated *renin* mRNA expression at early-, mid-, and late-stages of nephrogenesis (i.e., E14.5, E16.5, and E18.5) in control and *Tcf21* stromal cKO. Using in-situ hybridization (RNAscope), we detected a strong *Renin* signal in the arterioles leading into the developing glomeruli of control kidneys at E16.5 and E18.5 (**Fig.2 IA-B, IIA-D**). *Renin* expressing cells were primarily distributed in aggregates along the corticomedullary junction where newly formed glomeruli are known to mature (**Fig.2 IA**). In-line with the previously reported normal emergence of renin-expressing cells around E13.5 of the developing mouse kidney, we detected a scattered pattern of *Renin* expression at E14.5 of control kidneys which was less robust compared with *renin* expression at E16.5 and E18.5 (**Fig.2 IIE,F**). In contrast to controls, *Renin* mRNA expression in *Tcf21* stromal cKO kidneys was reduced, showing fewer *Renin*-positive areas, although cells expressing *renin* mRNA were still present in some pre-glomerular arterioles (**Fig.2 IA’**,**B’, IIA’-F’**).

**Figure 2:**
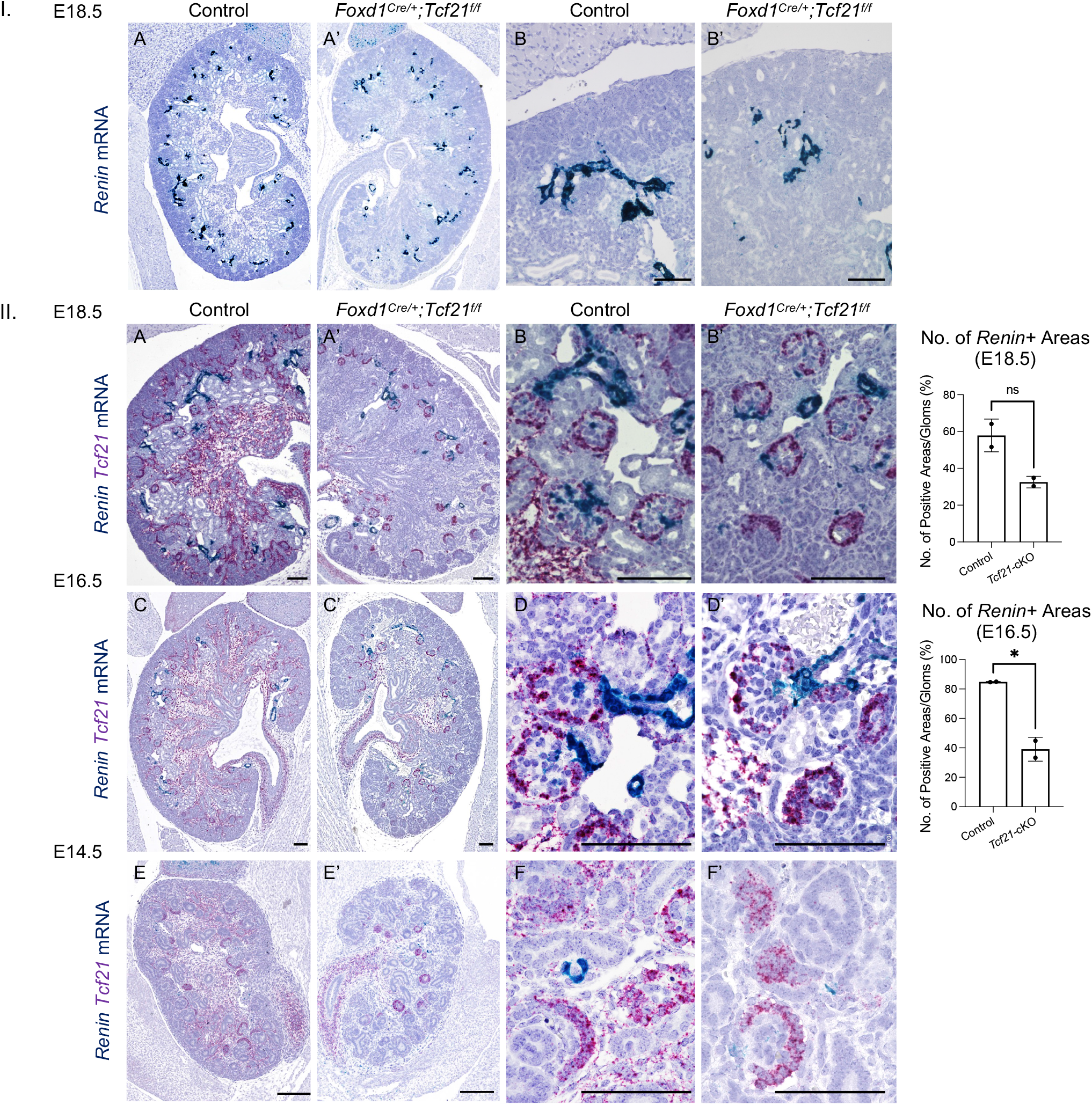
Reduced renin mRNA expression in the developing kidney of *Tcf21* stromal cKO. In-situ hybridization *(RNAscope)*. **I**. A-B. At E16.5, control kidneys express renin in cells of the afferent arterioles and in the hilum of the glomerulus along the corticomedullary junction, while *Foxd1*^*Cre/+*^*;Tcf21*^*f/f*^ kidneys demonstrate lower renin expression. *n*=4. **II**. A-F. Control kidneys at E14.5 through E18.5 demonstrate robust Tcf21 expression (purple) in the stroma and glomeruli, and renin expression (blue) along the afferent arterioles leading to the glomerulus hilum. A’-F’. *Foxd1*^*Cre/+*^*;Tcf21*^*f/f*^ kidneys show a marked reduction in Tcf21 expression in most of the stroma. However, Tcf21 expression remains intact in the glomeruli and stroma surrounding the ureter, where cells that originate from *Six2+* and *Tbx18+* progenitors, respectively. Renin expression is diminished in the JG regions of *Tcf21* stromal cKO. (*Right*) Quantifications of mRNA renin-positive areas (cell aggregates) in controls and *Tcf21* stromal cKO kidneys at E18.5 and E16.5, normalized to the total number of glomeruli (*p*=0.062 and *p*=0.031, respectively). *n*=4 at each timepoint. Sex not determined. Scale bar 100μm.

As to *Tcf21* expression in controls, as previously described (17), we detected robust *Tcf21* signal throughout stromal regions surrounding the cortex, sub-cortex, medulla, papilla, and ureter (**Fig.2 IIA,C,E**). In addition to the stroma, a strong Tcf21 signal was detected in podocytes at E16.5 and E18.5, and in segments of developing nephron structures at E14.5 (**Fig.2 IIB,D,F**). On the other hand, *Foxd1*^*Cre/+*^*;Tcf21*^*f/f*^ kidneys showed marked reduction in *Tcf21* expression in the stroma, but its expression was still detected in some regions of the medullary stroma (**Fig.2 IIC’**), which could be explained by incomplete Cre-mediated excision of *Tcf21* in a population of stromal cells. As expected, we detected *Tcf21* signal in podocytes, developing nephrons, and in the ureteric stromal of the cKOs, as these cells are not progeny of the Foxd1+ lineage but originate from the Six2 and Tbx18 lineages (22, 23). Taken together, these results suggest that expression of *Tcf21* in Foxd1+ cells is required for normal expression of *renin* mRNA and protein with ramifications to the development of the afferent arterioles.

### Inactivation of *Tcf21* under the *Renin* promotor does not recapitulate the vascular phenotype of *Foxd1*^***Cre/+***^***;Tcf21***^***f/f***^

The vascular defect seen in *Tcf21* stromal cKO kidneys brought up the possibility that continuous *Tcf21* expression in JG cells might be necessary for normal arterial development. To test this, we inactivated *Tcf21* in renin-expressing cells by breeding *Tcf21*^*f/f*^ mice with *Ren1*^*dCre/+*^ mice (14, 24, 25), creating the *Ren1*^*dCre/+*^*;Tcf21*^*f/f*^ (*Tcf21* renin cKO) mouse. Using immunostaining with the pan-endothelial marker PECAM (CD31) and 3D image analysis of cleared E16.5 kidneys, we observed no difference in vascular morphology between the *Tcf21* <u>renin</u> cKO and controls (**Fig.3 IA, A’, Suppl. Fig.1**). In contrast, *Tcf21* stromal cKO kidneys displayed fewer, shorter, and thinner branches (**Fig.3 IB-E** (17), quantification in **Fig.3 II**). This suggests that once Foxd1+ cells become renin cells, *Tcf21* expression is no longer needed for kidney vasculature formation. Evaluating the morphology of the *Ren1*^*dCre/+*^*;Tcf21*^*f/f*^ kidneys in two- and four-month-old mice showed normal renal architecture and vessel morphology compared with controls (**Fig.3 IIIA-H, Suppl. Fig.2**), indicating Tcf21 is dispensable once renin cell identity is established.

**Figure 3:**
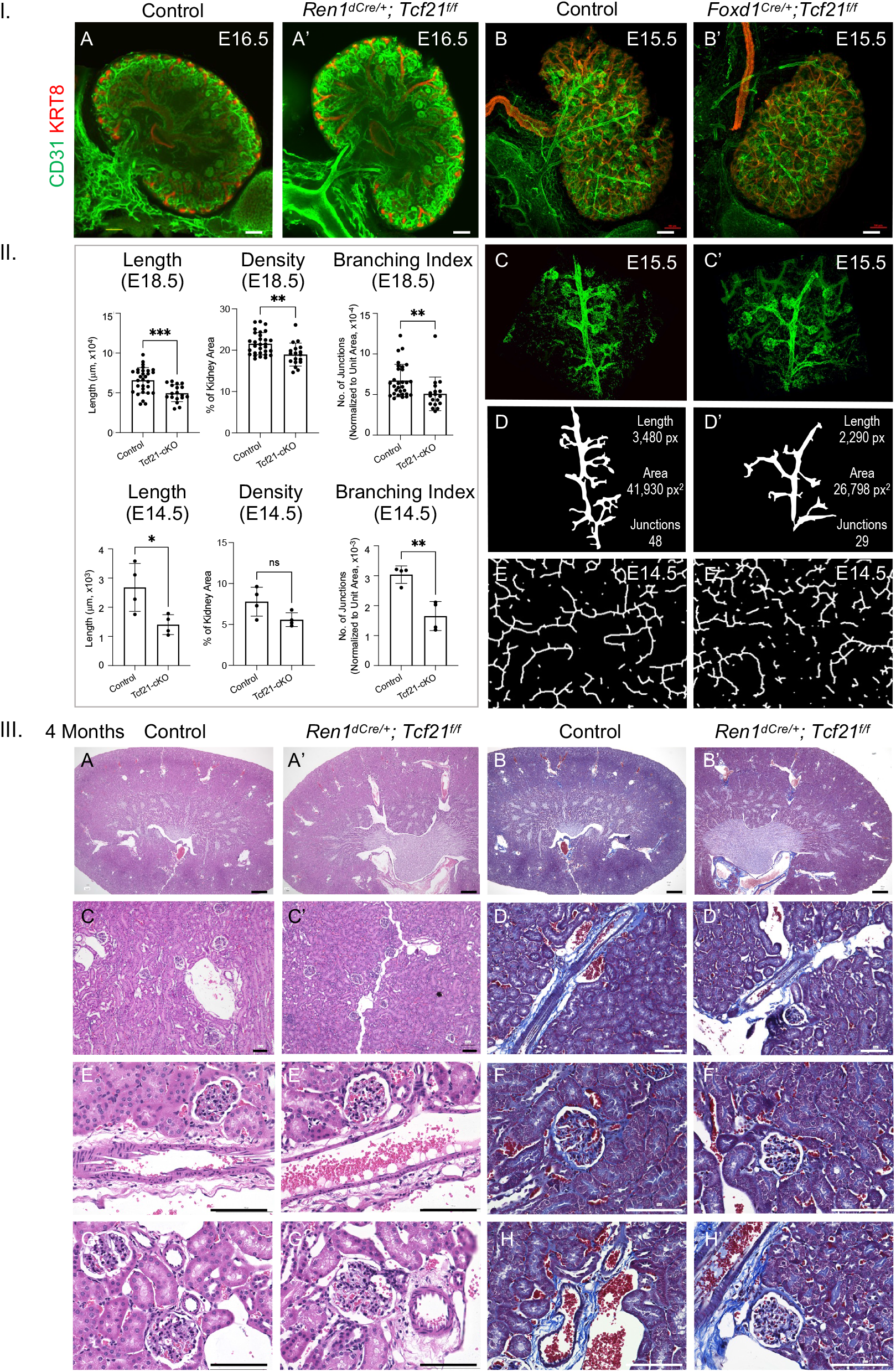
Inactivation of Tcf21 via the renin promoter does not recapitulate the vascular phenotype of *Foxd1*^*Cre/+*^*;Tcf21*^*f/f*^ kidneys. **I**. Immunostaining for CD31 (PECAM, pan-endothelial) and Keratin8 (ureteric bud) demonstrates the arterial tree of E16.5 *Ren1*^*dCre/+*^*;Tcf21*^*f/f*^ (left) and E15.5 *Foxd1*^*Cre/+*^*;Tcf21*^*f/f*^ (right) kidneys and controls. *n*=8, 6 males, 2 females. A. The *Ren1*^*dCre/+*^; *Tcf21*^*f/f*^ shows vascular patterning that is unchanged compared with control (3D visualization in **Suppl Fig. 1**). B. In contrast, the arterial tree of *Foxd1*^*Cre/+*^*;Tcf21*^*f/f*^ kidneys demonstrates a distorted pattern with disorganized, shorter, and thinner arterioles that display abnormal centripetal pattern with abnormal vessel branching of an aberrant ramification leading into the capsule (described in detail in (17)). C. Magnification of the *Foxd1*^*Cre/+*^*;Tcf21*^*f/f*^ arterial tree and D, vessel tracing. E. At E14.5, vessel tracing of CD31 in *Foxd1*^*Cre/+*^*;Tcf21*^*f/f*^ kidneys shows shortened and disorganized vasculature in the medullary region. *n*=8, 6 males, 2 females. **II**. Quantifications of vasculature in E18.5 *Foxd1*^*Cre/+*^*;Tcf21*^*f/f*^ kidneys (top): vessel length (*p*=0.0004), density (*p*=0.0023), and branching index (*p*=0.0100), and in E14.5 *Foxd1*^*Cre/+*^*;Tcf21*^*f/f*^ kidneys (bottom): vessel length (*p*=0.0284), density (*p*=0.0656), and branching index (*p*=0.0027) compared with controls. **III**. Histology of four-month-old *Ren1*^*dCre/+*^; *Tcf21*^*f/f*^ kidneys does not show phenotypic changes upon H&E (left) and Trichrome (right) staining compared with controls. At 4 months, similar to the findings at E15.5, the morphology of the vascular walls and glomeruli appear intact in *Ren1*^*dCre/+*^; *Tcf21*^*f/f*^. Scale bar 1000 μm for III. A-B, 100 μm for III. C-H. *n*=4, males.

To test if Tcf21 regulates renin expression, we compared *renin* positive spots in *Ren1*^*dCre/+*^*;Tcf21*^*f/f*^ and control kidneys using in-situ hybridization. In two-month-old mice, *Tcf21* renin cKO kidneys showed similar *renin* mRNA expression pattern as controls (**Suppl. Fig.3**). Similarly, at E14.5, *Tcf21* and *renin* mRNA signals did not co-localize (**Suppl. Fig.4**), suggesting *Tcf21*’s regulatory action is completed before *renin* promoter activation.

### Temporal Dynamics of Tcf21 Expression in the Differentiation of Foxd1+ Cells into Juxtaglomerular Cells

To investigate the role of Tcf21 in the differentiation of Foxd1+ cells into JG cells, we mapped the developmental trajectory from metanephric mesenchyme progenitors to mature renin-expressing JG cells. This was achieved through an integrated analysis of single-cell RNA sequencing (scRNA-seq) and single-cell ATAC sequencing (scATAC-seq) on purified GFP+ cells from *Foxd1*^*Cre/+*^*;Rosa26*^*mTmG*^ control mouse kidneys (*n=*6,191 cells), representing the stromal lineage. These analyses build upon the scRNA-seq data previously reported in (26), where we investigated renin cell development using chromatin accessibility and single-cell transcriptomics (GSE218570). In the current study, we specifically explore the temporal dynamics of Tcf21 in the JG cell lineage. These analyses were conducted at four key developmental stages: E12, E18, postnatal day 5 (P5), and postnatal day 30 (P30), capturing critical phases of kidney development. Clustering of the dataset identified a predominant population of stromal derivatives and a smaller proportion of non-stromal lineages such as ureteric bud cells and podocytes (**Suppl Fig. 5A-C**). To construct a pseudotime trajectory depicting the differentiation of Foxd1+ stromal progenitors to mature renin-expressing cells, we first identified cells expressing *Foxd1* at or above the third quartile and with a predicted gene score based on chromatin accessibility in the upper quartile. Next, we identified cells with high combined expression and chromatin accessibility scores for *Ren1* and *Akr1b7* in the upper quartile. Early developmental populations with elevated *Foxd1* levels were designated as the trajectory’s starting point, while late developmental cells with high Ren1/Akr1b7 expression served as endpoints. We utilized the addTrajectory() and plotTrajectory() functions from ArchR to visualize dynamic chromatin and gene expression changes, with a specific focus on the temporal dynamics of Tcf21 (**Suppl Fig. 5D-G**). This selection captured nine populations spanning the JG lineage, from metanephric mesenchyme to mature renin expressing cells (*n=*2,054) (**Fig. 4A**). These populations expressed markers indicative of stromal cells and vascular mural cells (including vascular smooth muscle cells, pericytes, renin cells) to varying degrees (**Fig. 4A**). The pseudotime analysis revealed developmental states within the JG lineage, mapping the progression from metanephric mesenchyme to fully differentiated JG cells (**Fig. 4B**).

**Figure 4:**
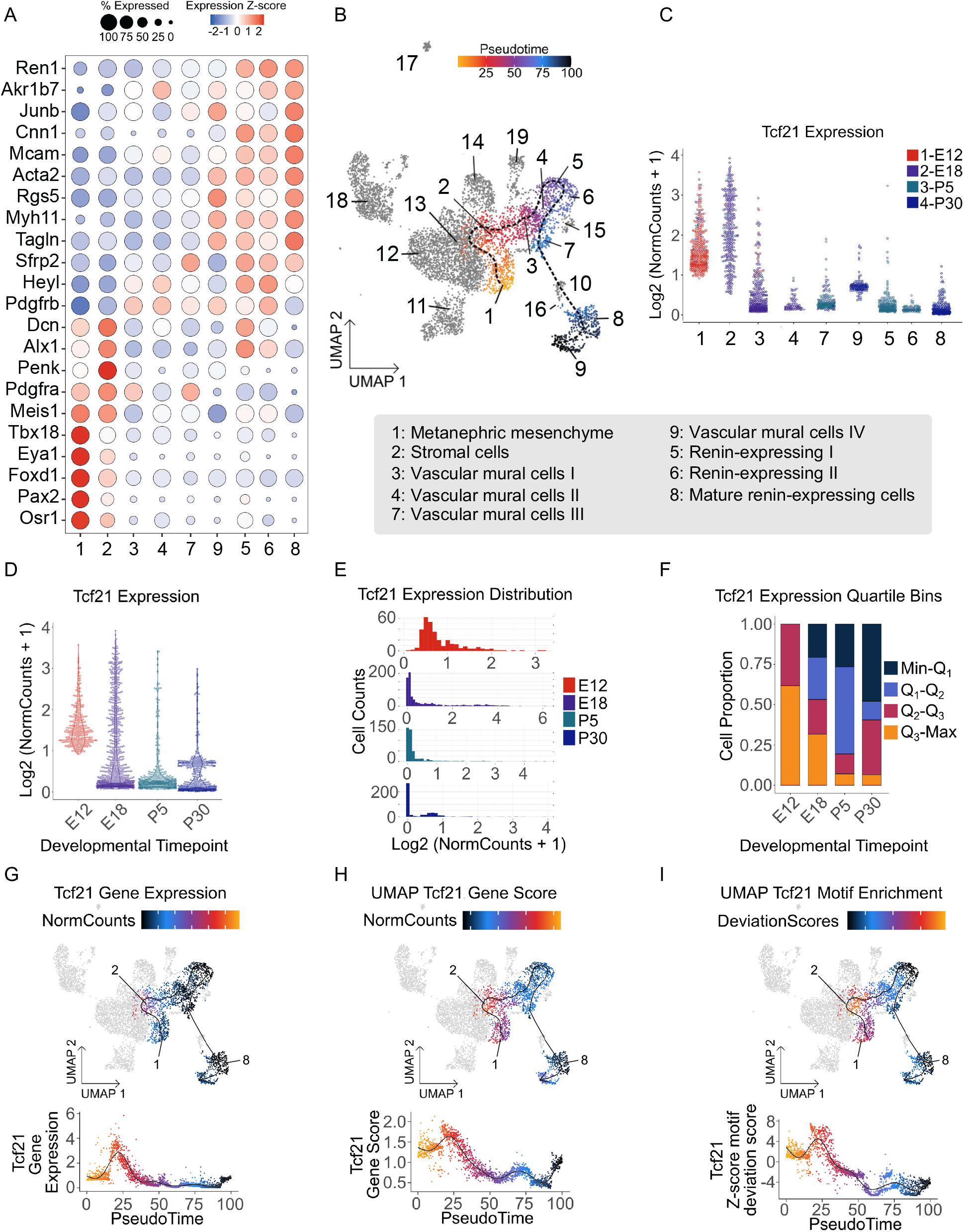
Distribution of Tcf21 expression from integrated scRNA-seq and scATAC-seq data analysis. A. Bubble plot depicting the expression of key cell type markers across the populations of the JG lineage (combined for E12, E18, P5, and P30) (see details in Suppl Fig. 5), illustrating the transition through various developmental stages. B. Uniform Manifold Approximation and Projection (UMAP) visualization of the pseudotime trajectory mapping the differentiation from metanephric mesenchyme progenitors to mature renin-expressing JG cells (*n*=2,054 cells), shown by cell identity. A scale bar represents the progression of pseudotime, with cells (dots) positioned closer to the start of the trajectory being at earlier state of differentiation compared to cells positioned later in pseudotime. C. Violin plot of Tcf21 expression across all JG lineage clusters and developmental timepoints. D. Violin plot showing Tcf21 expression levels (log2 scale) across developmental time points. E. Histogram presenting the number of JG cells across different Tcf21 expression levels (x-axis) divided by developmental time points. F. Tcf21 expression presented in quartile bins across developmental time points. Note the highest Tcf21 expression at E12, with lower expression levels observed at later stages of kidney development (E18, P5) and in the mature kidney (P30). G-H. (Upper plots) Tcf21 RNA expression levels within the JG lineage presented as signal intensity on a log2 scale in G and inferred gene expression from chromatin accessibility (Gene Score) on a log2 scale in H projected on the corresponding UMAPs. (Lower plots) Corresponding values plotted along the pseudotime of the JG cell trajectory. Note the higher Tcf21 expression in clusters identified as metanephric mesenchyme (cluster 1) and stromal cells (cluster 2), while Tcf21 expression declines as cell progress to mature renin-expressing cells (cluster 8). I. (Upper plot) Tcf21 motif enrichment presented as z-scores of the bias-corrected motif deviation values within the JG lineage projected on the UMAP. (Lower plots) Corresponding values plotted along the pseudotime. Number of animals: *n*=15 (E12), *n*=18 (E18), *n*=12 (P5), and *n*=10 (P30). Both sexes represented based on Mendelian ratios.

A violin plot of *Tcf21* expression levels in [log2 (normcounts + 1)] across these populations revealed high expression of Tcf21 in cells of the metanephric mesenchyme (cluster 1) and stromal cells (cluster 2), with decreasing levels as stromal cells differentiate into mature renin-expressing cells (cluster 8) (**Fig. 4C**). A similar pattern is evident in a violin plot of Tcf21 expression across the four key developmental timepoints in the JG lineage, demonstrating a decline in expression from E12 to P30 (**Fig. 4D**). Additionally, a histogram depicting the distribution of *Tcf21* expression split by developmental stages supports this, showing that most E12 cells express higher levels of *Tcf21* compared to later stages (**Fig. 4E**). Quartile analysis further confirmed this trend, with a higher proportion of cells expressing Tcf21 at or above the second quartile at E12 compared to later stages (**Fig. 4F**). Taken together, this data suggest that *Tcf21* is more prevalent during early developmental stages (E12), supporting its critical role in at initial differentiation of progenitor cells.

Motif enrichment and pseudotime trajectory analyses further emphasize Tcf21’s role in early kidney development (**Fig. 4G-I**), consistent with findings from the morphological immunostaining analysis described above.

Overall, these results indicate that Tcf21 is highly active during early stages of kidney development, particularly in metanephric mesenchyme and stromal progenitors, with its influence in differentiating JG cells diminishing as development proceeds.

## DISCUSSION

The developmental pathways that guide the differentiation of Foxd1+ stromal progenitors into various cell types, such as JG cells, are crucial for proper kidney morphogenesis and function. Our study elucidates the role of Tcf21 in the differentiation of Foxd1+ progenitor cells into JG cells during kidney development. The results demonstrate that Tcf21 expression is essential for the early specification of these progenitors into renin-expressing JG cells, with its expression peaking during early embryogenesis and diminishing as development progresses. The inactivation of Tcf21 in Foxd1+ cells led to a significant reduction in renin-positive areas within the developing kidney. This was evident through the decreased renin protein and mRNA expression, as well as the morphological alterations in the afferent arterioles. Our results suggest that Tcf21 primes Foxd1+ progenitor cells for differentiation into the JG lineage, supporting its role as a “founder transcription factor.” Founder factors like Tcf21 initiate early lineage specification, guiding progenitor cells toward specialized fates. The reduced renin and MCAM expression in the *Tcf21* stromal cKO model suggests that Tcf21 plays a critical role in the proper differentiation and function of renin-expressing pericytes, which are important for vessel formation and renin synthesis. Our findings align with previous research indicating the importance of Foxd1+ progenitors in kidney morphogenesis and vascular development. The temporal dynamics of Tcf21 expression and motif enrichment, revealed through integrated scRNA-seq and scATAC-seq analysis, underscore its critical role in the early stages of JG lineage specification. The observed decline in Tcf21 as metanephric mesenchyme progenitors and stromal cells differentiate into renin-expressing for vascular development. The normal renal architecture and vessel morphology in mature *Tcf21* renin cKO cells highlights its essential involvement in the initial phases of JG cell development. Consistent with this observation is the finding that inactivation of Tcf21 under the renin promoter did not replicate the vascular phenotype observed in the *Foxd1*^*Cre/+*^*;Tcf21*^*f/f*^ model. This suggests that once Foxd1+ cells commit to the renin-expressing lineage, Tcf21 is no longer required kidneys further support this notion, suggesting that Tcf21’s role is primarily confined to the early stages of cell fate determination. These findings provide new insights into the molecular mechanisms governing JG cell differentiation and emphasize the role of Tcf21 in the early specification of Foxd1+ cells into JG cells.

It is important to mention that the variability in Tcf21 deletion among cells, which may have resulted from non-absolute Cre-mediated recombination, possibly affected the uniformity of our findings. Additionally, this study did not address the complexity of the transcriptional networks governing Foxd1+ cell differentiation in terms of identifying Tcf21 binding partners, coregulators, and direct targets. These aspects should be explored in future research and could have functional implications. Finally, we did not explore potential compensatory mechanisms that might mitigate the effects of Tcf21 deficiency on JG cell differentiation. Although we did not consistently determine the sex of the mice in every experiment, and there was variability in the representation of each sex across different experiments, both preliminary and subsequent analyses consistently demonstrated a vascular phenotype in both male and female *Foxd1*^*Cre/+*^*;Tcf21*^*f/f*^ and the absence of one in *Ren1*^*dCre/+*^*;Tcf21*^*f/f*^ mice. To account for potential sex-related effects, we utilized age-matched male and female cohorts, selecting them based on availability at the time of each experiment. Furthermore, our preliminary data and additional experiments, not included in this manuscript, did not reveal any significant gross morphological differences between the sexes.

In summary, our study highlights the complexity of transcriptional regulation in the specification and maintenance of the JG cell lineage. Tcf21 may prime cis-regulatory elements in a cell- and developmental stage-specific manner to control the specification of various stromal sub-populations. Further work is required to construct a comprehensive spatial and temporal developmental atlas of stromal cells during kidney development. By uncovering Tcf21’s role in early specification and its dispensability in later stages, we lay a foundation for further research into key signaling pathways and mechanisms in renal vascular development.

## Supporting information

3D imaging of control kidney

3D imaging of mutant kidney

## DATA AVAILABILITY

Data is publicly available at Gene Expression Omnibus (GSE218570).

## SUPPLEMENTAL MATERIAL

Supplemental Figures 1-5:

Suppl Fig. 1: Tissue clearing and immunostaining of *Ren1*^*dCre/+*^*;Tcf21*^*f/f*^. n=4, females.

Suppl Fig. 2: Histology of two-month old mice of control and *Ren1*^*dCre/+*^*;Tcf21*^*f/f*^ kidneys. n=16, females.

Suppl Fig. 3: Normal renin mRNA expression in *Ren1*^*dCre/+*^*;Tcf21*^*f/f*^ mouse kidney at 2 months.

Suppl Fig. 4: RNAScope of *Tcf21* and *renin* at E14.5 control kidneys. n=2, unknown sex.

Suppl Fig. 5: Juxtaglomerular (JG) Cell Psuedotime Trajectory via scRNA-seq and scATAC-seq.

Supplemental Table 1:

Suppl Table 1: Primer information (genotyping)

## ACKNOWLEDGMENTS

Confocal microscopy was provided by the Center for Advanced Microscopy of Northwestern University and by the Microscopy and Histology Group at Stanley Manne Children’s Research Institute affiliated with Ann and Robert H. Lurie Children’s Hospital of Chicago. With gratitude to the Zell Family Foundation.

## Author contributions

G.F., M.L.S.S.L., R.A.G., A.G.M., and S.E.Q. designed the study; G.Y., G.F., H.A., and A.G.M., carried out experiments; G.Y., G.F., M.L.S.S.L., R.A.G., J.P.S., and H.A., analyzed the data; G.F., and G.Y., made the figures; G.F., M.L.S.S.L., R.A.G., J.P.S., G.Y., and S.E.Q. drafted and revised the paper; all authors approved the final version of the manuscript.

## LEGENDS FOR SUPPLAMENTRAY FIGURE

**Suppl Figure 1: Tissue clearing and immunostaining of kidneys from *Ren1***^***dCre/+***^***;Tcf21***^***f/f***^ **and controls**. Babb clearing, immunostaining with the pan-endothelial marker PECAM (CD31; red), and 3D imaging show no difference in vascular morphology between *Ren1*^*dCre/+*^*;Tcf21*^*f/f*^ and control kidneys at E16.5. *n*=4, females.

**Suppl Figure 2: Histology from 2-month-old *Ren1***^***dCre/+***^***;Tcf21***^***f/f***^ **and control kidneys**. H&E (top) and Trichome (bottom) stainings reveal no gross phenotypic changes in *Ren1*^*dCre/+*^*;Tcf21*^*f/f*^ kidneys compared with controls. Vascular wall and glomeruli morphology appear intact. *n*=16, females. Scale bar 100 μm.

**Suppl Figure 3**: **Normal *renin* mRNA expression in *Ren1***^***dCre/+***^***;Tcf21***^***f/f***^ **kidneys at 2 months**.

In-situ hybridization *(RNAscope)*. A-B. *Renin* expression (blue) is detected in the arteriolar walls (yellow arrows) and in the glomerulus hilum (black arrows) similarly in 2-month-old *Ren1*^*dCre/+*^; *Tcf21*^*f/f*^ and control kidneys. C. The distribution of renin-expressing cells is similar in *Ren1*^*dCre/+*^; *Tcf21*^*f/f*^ and controls. *n*=4, females. Scale bar 75μm (A), 100μm (B,C).

**Suppl Figure 4: In-situ hybridization of Tcf21 and renin in E14.5 *Ren1***^***d+/+***^***;Tcf21***^***f/f***^ **kidneys**. In-situ hybridization (*RNAscope*) shows no co-localization of *Tcf21* (purple) and *renin* (blue) mRNA in E14.5 control kidneys. This time point was chosen because renin-expressing cells emerge around E13.5. *n*=2, sex not determined.

**Suppl Figure 5**: **Juxtaglomerular (JG) Cell Psuedotime Trajectory via scRNA-seq and scATAC-seq**. A. UMAP visualization showing clustering of the entire GFP+ cell population isolated from *Foxd1*^*Cre/+*^*;Rosa26*^*mTmG/+*^ control mouse kidneys at E12, E18, P5, and P30 (*n=*6,191 cells). A total of 19 populations were identified. Populations representing stromal derivatives (clusters 2-16) are highlighted in red. B. Bubble plot demonstrating expression of key kidney cell lineage markers across the 19 clusters. C. Cluster heatmap showing marker expression across all GFP+ clusters. D. UMAP visualization of the pseudotime trajectory mapping the differentiation form metanephric mesenchyme progenitors to mature renin-expressing JG cells (*n=*2,054 cells), shown by embryonic stage. Early populations with high *Foxd1* expression serve as the starting point, and late populations with high *Ren1/Akr1b7* expression were identified as endpoints. E-G. Feature plots visualizing Foxd1, Renin, and Akr1b7 expression levels across the entire GFP+ cell populations.

## REFERENCES

1. Sequeira-Lopez MLS, and Gomez RA. Renin Cells, the Kidney, and Hypertension. Circ Res 128: 887–907, 2021.

2. Sequeira Lopez ML, and Gomez RA. Development of the renal arterioles. J Am Soc Nephrol 22: 2156–2165, 2011.

3. Reddi V, Zaglul A, Pentz ES, and Gomez RA. Renin-expressing cells are associated with branching of the developing kidney vasculature. J Am Soc Nephrol 9: 63–71, 1998.

4. Gomez RA, Pentz ES, Jin X, Cordaillat M, and Sequeira Lopez ML. CBP and p300 are essential for renin cell identity and morphological integrity of the kidney. Am J Physiol Heart Circ Physiol 296: H1255–1262, 2009.

5. Smith DL, Morris BJ, Do YS, Law RE, Shaw KJ, and Hseuh WA. Identification of cyclic AMP response element in the human renin gene. Biochem Biophys Res Commun 200: 320–329, 1994.

6. Pan L, Black TA, Shi Q, Jones CA, Petrovic N, Loudon J, Kane C, Sigmund CD, and Gross KW. Critical roles of a cyclic AMP responsive element and an E-box in regulation of mouse renin gene expression. J Biol Chem 276: 45530–45538, 2001.

7. Yamaguchi H, Gomez RA, and Sequeira-Lopez MLS. Renin Cells, From Vascular Development to Blood Pressure Sensing. Hypertension 80: 1580–1589, 2023.

8. Tandon P, Miteva YV, Kuchenbrod LM, Cristea IM, and Conlon FL. Tcf21 regulates the specification and maturation of proepicardial cells. Development 140: 2409–2421, 2013.

9. Acharya A, Baek ST, Huang G, Eskiocak B, Goetsch S, Sung CY, Banfi S, Sauer MF, Olsen GS, Duffield JS, Olson EN, and Tallquist MD. The bHLH transcription factor Tcf21 is required for lineage-specific EMT of cardiac fibroblast progenitors. Development 139: 2139–2149, 2012.

10. Sequeira-Lopez ML, Lin EE, Li M, Hu Y, Sigmund CD, and Gomez RA. The earliest metanephric arteriolar progenitors and their role in kidney vascular development. Am J Physiol Regul Integr Comp Physiol 308: R138–149, 2015.

11. Song R, Lopez M, and Yosypiv IV. Foxd1 is an upstream regulator of the renin-angiotensin system during metanephric kidney development. Pediatr Res 82: 855–862, 2017.

12. Kragesteen BK, Giladi A, David E, Halevi S, Geirsdottir L, Lempke OM, Li B, Bapst AM, Xie K, Katzenelenbogen Y, Dahl SL, Sheban F, Gurevich-Shapiro A, Zada M, Phan TS, Avellino R, Wang SY, Barboy O, Shlomi-Loubaton S, Winning S, Markwerth PP, Dekalo S, Keren-Shaul H, Kedmi M, Sikora M, Fandrey J, Korneliussen TS, Prchal JT, Rosenzweig B, Yutkin V, Racimo F, Willerslev E, Gur C, Wenger RH, and Amit I. The transcriptional and regulatory identity of erythropoietin producing cells. Nat Med 29: 1191–1200, 2023.

13. Maezawa Y, Onay T, Scott RP, Keir LS, Dimke H, Li C, Eremina V, Maezawa Y, Jeansson M, Shan J, Binnie M, Lewin M, Ghosh A, Miner JH, Vainio SJ, and Quaggin SE. Loss of the podocyte-expressed transcription factor Tcf21/Pod1 results in podocyte differentiation defects and FSGS. J Am Soc Nephrol 25: 2459–2470, 2014.

14. Mohamed TH, Watanabe H, Kaur R, Belyea BC, Walker PD, Gomez RA, and Sequeira-Lopez MLS. Renin-Expressing Cells Require beta1-Integrin for Survival and for Development and Maintenance of the Renal Vasculature. Hypertension 76: 458–467, 2020.

15. Kobayashi A, Mugford JW, Krautzberger AM, Naiman N, Liao J, and McMahon AP. Identification of a multipotent self-renewing stromal progenitor population during mammalian kidney organogenesis. Stem Cell Reports 3: 650–662, 2014.

16. Wang F, Flanagan J, Su N, Wang LC, Bui S, Nielson A, Wu X, Vo HT, Ma XJ, and Luo Y. RNAscope: a novel in situ RNA analysis platform for formalin-fixed, paraffin-embedded tissues. J Mol Diagn 14: 22–29, 2012.

17. Finer G, Maezawa Y, Ide S, Onay T, Souma T, Scott R, Liang X, Zhao X, Gadhvi G, Winter DR, Quaggin SE, and Hayashida T. Stromal Transcription Factor 21 Regulates Development of the Renal Stroma via Interaction with Wnt/beta-Catenin Signaling. Kidney360 3: 1228–1241, 2022.

18. Chen J, Luo Y, Hui H, Cai T, Huang H, Yang F, Feng J, Zhang J, and Yan X. CD146 coordinates brain endothelial cell-pericyte communication for blood-brain barrier development. Proc Natl Acad Sci U S A 114: E7622–E7631, 2017.

19. Manocha E, Consonni A, Baggi F, Ciusani E, Cocce V, Paino F, Tremolada C, Caruso A, and Alessandri G. CD146(+) Pericytes Subset Isolated from Human Micro-Fragmented Fat Tissue Display a Strong Interaction with Endothelial Cells: A Potential Cell Target for Therapeutic Angiogenesis. Int J Mol Sci 23: 2022.

20. Espagnolle N, Guilloton F, Deschaseaux F, Gadelorge M, Sensebe L, and Bourin P. CD146 expression on mesenchymal stem cells is associated with their vascular smooth muscle commitment. J Cell Mol Med 18: 104–114, 2014.

21. Castrop H, Hocherl K, Kurtz A, Schweda F, Todorov V, and Wagner C. Physiology of kidney renin. Physiol Rev 90: 607–673, 2010.

22. Kobayashi A, Valerius MT, Mugford JW, Carroll TJ, Self M, Oliver G, and McMahon AP. Six2 defines and regulates a multipotent self-renewing nephron progenitor population throughout mammalian kidney development. Cell Stem Cell 3: 169–181, 2008.

23. Bohnenpoll T, Bettenhausen E, Weiss AC, Foik AB, Trowe MO, Blank P, Airik R, and Kispert A. Tbx18 expression demarcates multipotent precursor populations in the developing urogenital system but is exclusively required within the ureteric mesenchymal lineage to suppress a renal stromal fate. Dev Biol 380: 25–36, 2013.

24. Belyea BC, Xu F, Pentz ES, Medrano S, Li M, Hu Y, Turner S, Legallo R, Jones CA, Tario JD, Liang P, Gross KW, Sequeira-Lopez ML, and Gomez RA. Identification of renin progenitors in the mouse bone marrow that give rise to B-cell leukaemia. Nat Commun 5: 3273, 2014.

25. Sequeira Lopez ML, Pentz ES, Nomasa T, Smithies O, and Gomez RA. Renin cells are precursors for multiple cell types that switch to the renin phenotype when homeostasis is threatened. Dev Cell 6: 719–728, 2004.

26. Martini AG, Smith JP, Medrano S, Finer G, Sheffield NC, Sequeira-Lopez MLS, and Gomez RA. Renin Cell Development: Insights From Chromatin Accessibility and Single-Cell Transcriptomics. Circ Res 133: 369–371, 2023.

