## Supplementary material for "Tcf21 as a Founder Transcription Factor in Specifying Foxd1 Cells to the Juxtaglomerular Cell Lineage": Suppl Table and Figures

| Gene | Forward | Reverse |
|---|---|---|
| Tcf21$^{flox}$ | GTG TGC ATT TCT GTG GTT GTC TCT | CTG TTG TTT GTG CAG GTG GAG A |
| Foxd1$^{WT}$ | CTC CTC CGT GTC CTC GTC | TCT GGT CCA AGA ATC CGA AG |
| Foxd1$^{Cre}$ | GGG AGG ATT GGG AAG ACA AT | TCT GGT CCA AGA ATC CGA AG |
| Ren1d$^{WT}$ | GAA GGA GAG CAA AAG GTA AGA G | GTA GTA GAA GGG GGA GTT GTG |
| Ren1d$^{Cre}$ | CAC AGG CCC TGG GGT AAT AAA TCA AAG | CAG GCA AAT TTT GGT GTA CGG |
| Rosa26$^{WT}$ | CTC TGC TGC CTC CTG GCT TCT | CGA GGC GGA TCA CAA GCA ATA |
| Rosa26$^{mTmG}$ | CTC TGC TGC CTC CTG GCT TCT | TCA ATG GGC GGG GGT CGT T |
| Rbm31 (Sex) | CAC CTT AAG AAC AAG CCA ATA CA | GGC TTG TCC TGA AAA CAT TTG G |

**Supplemental Table 1: Primer information - genotyping**

3D – Mp4 file

**Supplemental Figure 1**

2-month-old

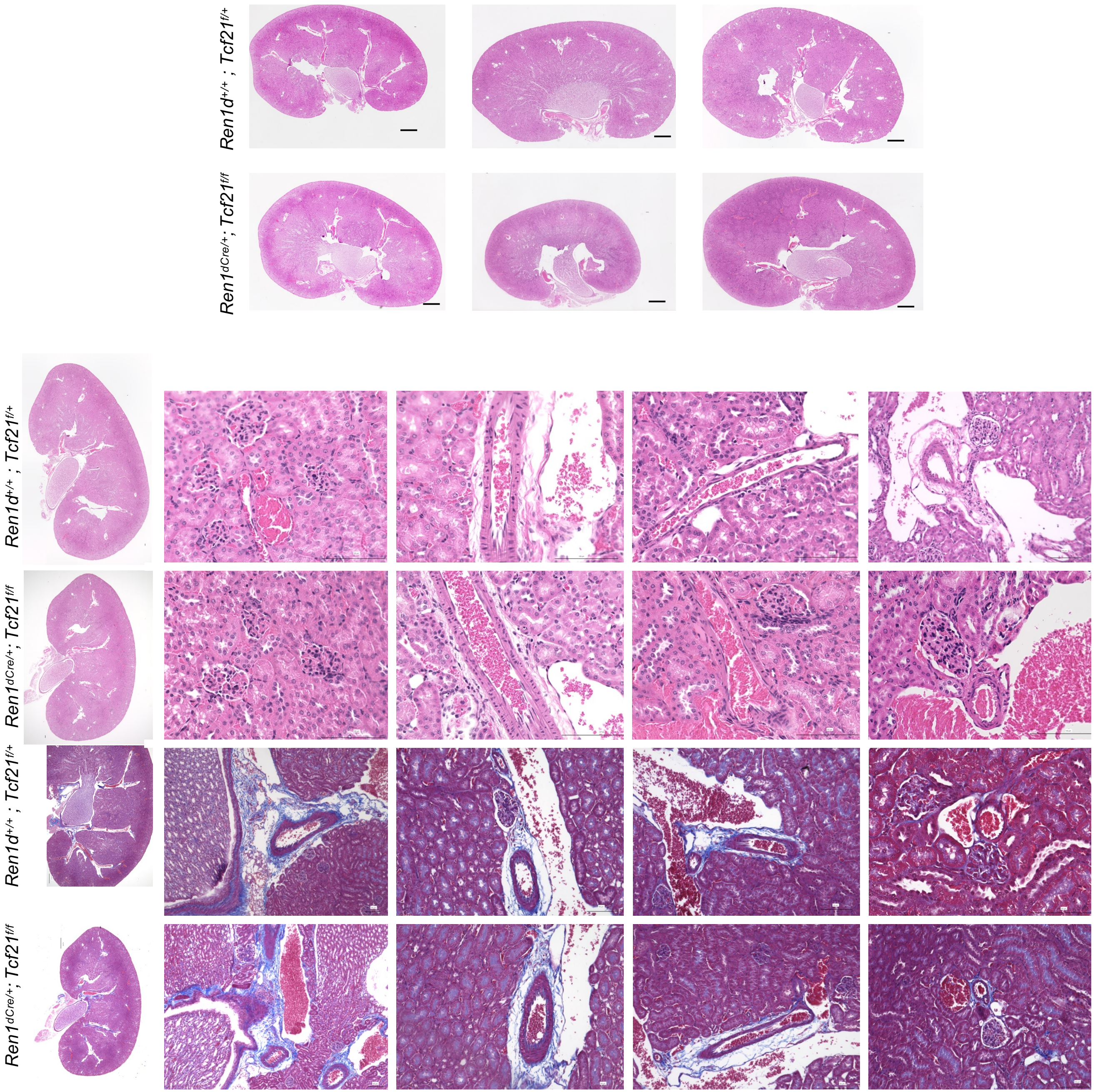


**Supplemental Figure 2**

2-month-old

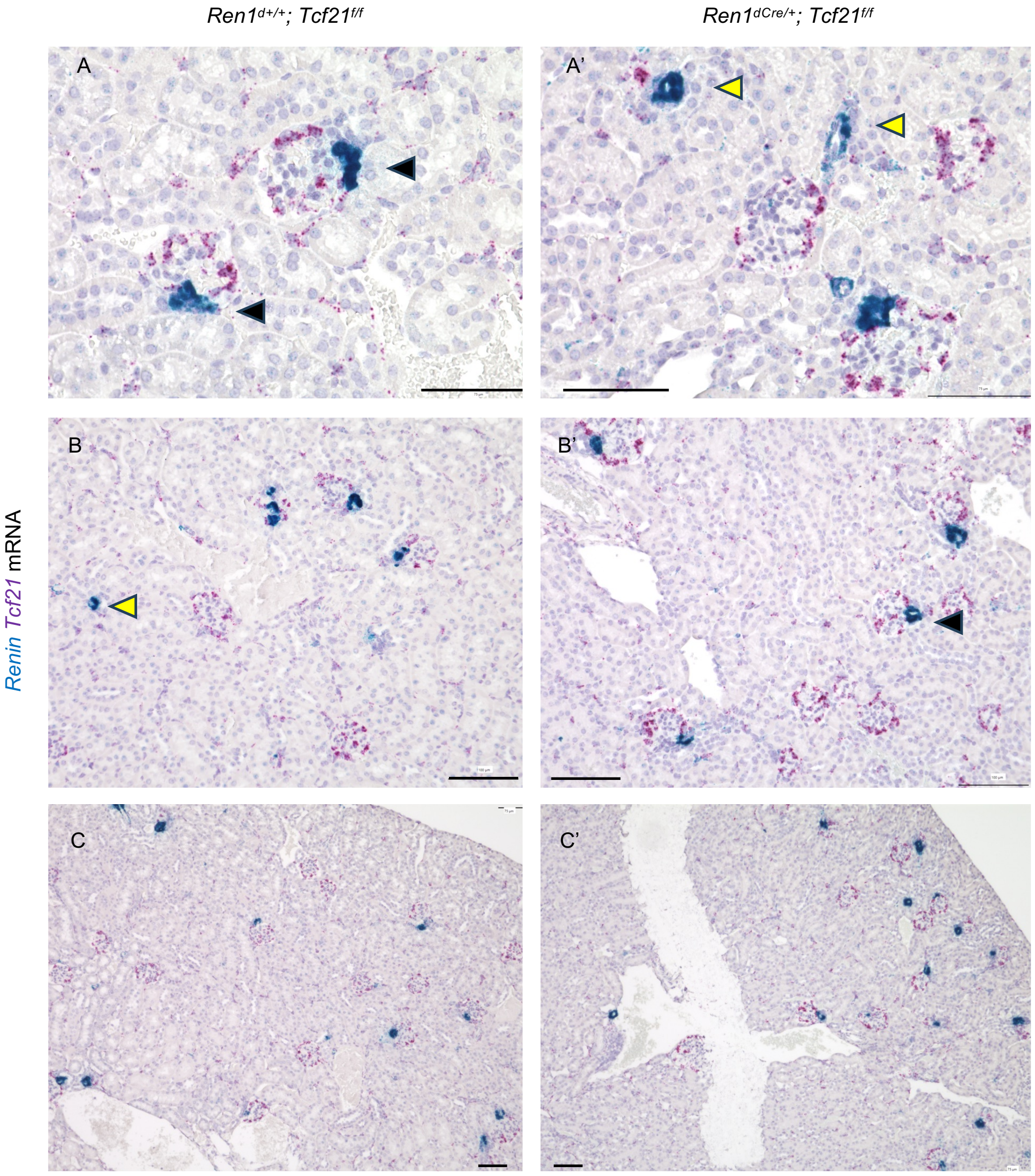


**Supplemental Figure 3**

E14.5 *Ren1*$^{d+/+}$; *Tcf21*$^{f/f}$

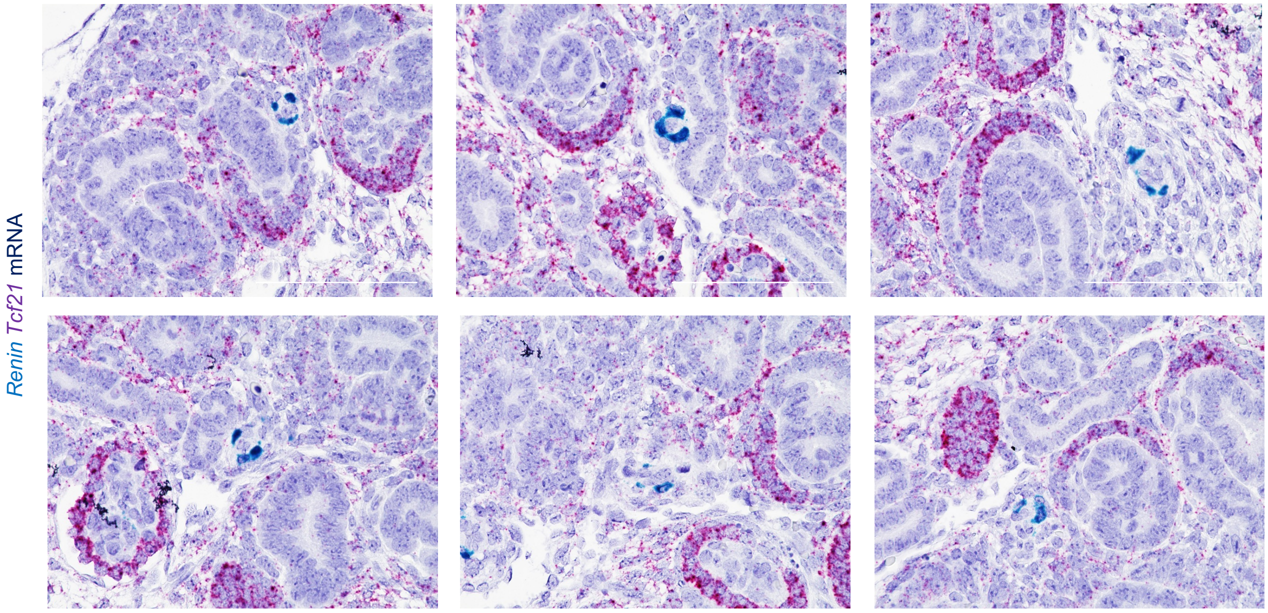


**Supplemental Figure 4**

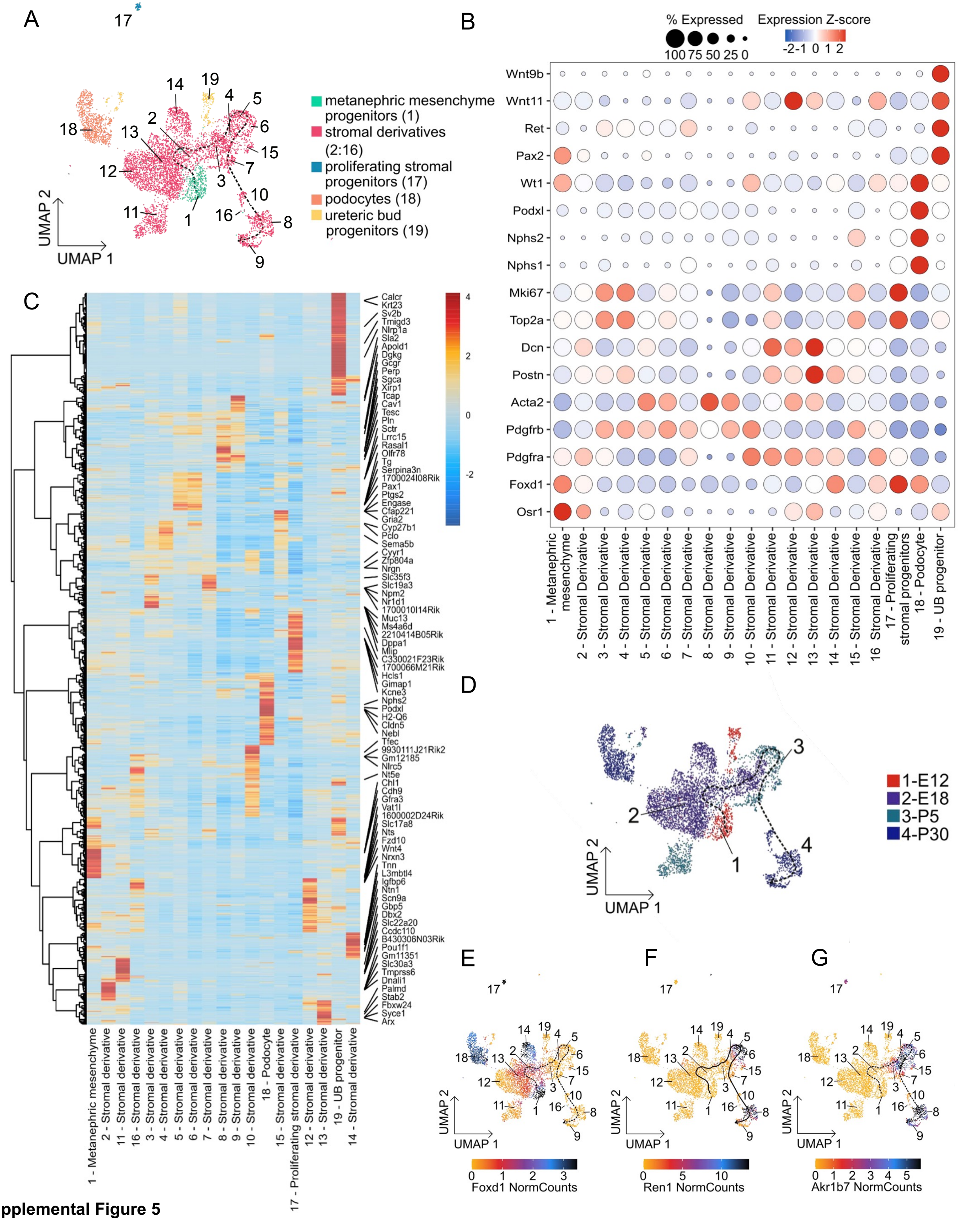


**Supplemental Figure 5**
